# Three-dimensional motion perception: comparing speed and speed change discrimination for looming stimuli

**DOI:** 10.1101/2020.04.03.023879

**Authors:** Abigail R. I. Lee, Justin M. Ales, Julie M. Harris

## Abstract

Judging the speed of objects moving in three dimensions is important in our everyday lives, because we interact with objects in a three-dimensional world. However, speed perception has been seldom studied for motion in depth, particularly when using monocular cues such as looming. Here, we compared speed discrimination, and speed change discrimination, for looming stimuli, to better understand what visual information is used for these tasks. For the speed discrimination task, we manipulated the distance and duration information available, to investigate if participants were specifically using speed information. For speed change discrimination, total distance and duration were held constant, hence they could not be used to successfully perform that task. We found speed change discrimination thresholds were consistently higher than those for speed discrimination. Evidence suggested that participants used a variety of cues to complete the speed discrimination task, not always solely relying on speed. Further, our data suggested that participants may switch between cues on a trial to trial basis. We conclude that speed change discrimination for looming is more difficult than speed discrimination, and that naїve participants may not always exclusively use speed for speed discrimination.

## Introduction

Perceiving the speed of objects moving towards us in the world is important in our daily lives, for example when safely crossing a road. Of particular importance is the ability to judge the speed, and speed changes, of objects approaching in three dimensions. There are both monocular and binocular sources of visual information we can use to judge these movements. Speed discrimination using binocular cues to motion in depth has been well studied (Brooks, 2002; Brooks & Mather, 2000; Brooks & Stone, 2004, 2006; Harris & Watamaniuk, 1995, 1996; Wardle & Alais, 2013). Perhaps more overlooked recently is the contribution of monocular cues to motion in depth, such as looming. Looming is usually defined as the change in retinal size that occurs when an object moves towards or away from an observer (e.g. Sekuler, 1992). The first evidence for the existence of mechanisms specifically sensitive to such change in size, that could be used for the perception of motion-in-depth, came from motion adaptation studies demonstrating that adaptation to size-change was separable to that for lateral motion (Beverley & Regan, 1979; Regan & Beverley, 1978).

In this study we investigate speed and speed change discrimination for looming stimuli, and we explore the strategies that naїve participants may be using for speed discrimination. We define looming as the expansion of the image of an object on the retina as it approaches, while the object in the world remains a constant size. When an object approaches or moves away from an observer at a constant speed in the world, the image of that object on the retina accelerates or decelerates respectively. The closer the object gets to the eye, the greater the acceleration. Therefore, to emulate real-world motion, our looming stimuli moved at a constant world speed, which resulted in an accelerating retinal speed (see Lee, Ales, & Harris, 2019).

Looming is thought to play a role in judging the speed of objects moving towards us. Speed discrimination for looming can be as sensitive as that for 2D motion, and is superior to that using other 3D motion cues. Speed discrimination thresholds for looming stimuli can be as low as 5% (Sekuler, 1992) similar to those for 2D motion (de Bruyn & Orban, 1988; Heidenreich & Turano, 1996; McKee, 1981; McKee, Silverman, & Nakayama, 1986; McKee & Welch, 1985; Orban, de Wolf, & Maes, 1984; Snowden & Braddick, 1991). By comparison, speed discrimination thresholds when using binocular cues to motion in depth are often much higher than that reported for looming and for 2D motion stimuli (Brooks & Stone, 2004, 2006, Harris & Watamaniuk, 1995, 1996). The higher sensitivity for looming cues over binocular cues suggests that looming is a critical cue for 3D motion perception.

However, speed discrimination tasks can be problematic to interpret. In traditional speed discrimination designs, it is impossible to be certain that participants judge speed, rather than distance or duration. Typically, if distance is kept constant, a participant could use speed or duration to make judgements. Conversely, if duration is held constant, participants can use speed or distance to make their judgements (for a review see McKee & Watamaniuk, 1994). To avoid this problem, speed change discrimination tasks have been developed (Monen & Brenner, 1994; Snowden & Braddick, 1991). In these tasks, observers are asked to discriminate a change in speed occurring during one interval, allowing total duration and distance to be held constant. In the standard interval, a stimulus travels at a constant speed. In the test interval, the stimulus travels slower and then an equal amount faster than the standard interval speed. Thus, the mean speed, and therefore the duration and distance, are kept constant whilst speed is varied. There is evidence from studies that explore 2D motion, and several types of 3D motion, that speed change discrimination is much more difficult than speed discrimination (e.g. Lee et al., 2019; Monen & Brenner, 1994; Snowden & Braddick, 1991).

No study has previously explored whether speed change discrimination for the monocular 3D motion cue of looming is more difficult than speed discrimination, and if so, why this may be. However, it is possible that speed discrimination may be an easier task because participants are able to use additional distance or duration cues to give their responses. We therefore had two aims:

1. To determine if speed change discrimination for looming is a more difficult task than speed discrimination.
2. To determine if participants use distance or duration information, rather than speed information, when it is available in speed discrimination tasks, and if this could explain the apparent difficulty of speed change discrimination judgements where these cues cannot be used.

## Methods

### Participants

Participants were required to have normal or corrected-to-normal vision and a stereoacuity of at least 120 arcseconds, as measured by the TNO test (16th edition). For the speed change discrimination task, 15 participants were recruited. Two participants had a stereoacuity of over 120 arcseconds, 1 participant was unable to do the task, and 1 participant stopped attending the testing sessions, leaving 11 participants who completed the experiment (8 female, 3 male, aged between 18-34). For the speed discrimination tasks, 9 new naive participants were recruited, as the participants from the speed change discrimination task could not be recalled in line with our ethical approval requirements. One participant from this new group had a stereoacuity of over 120 arcseconds, whilst another participant did not pass the training, leaving 7 participants who completed the experiment (5 female, 2 male, aged between 18-28). Participants gave informed consent before beginning the experiment and all procedures were approved by the University of St Andrews University Teaching and Research Ethics Committee (UTREC; Approval code: PS11904). All experiments adhered to the tenets of the Declaration of Helsinki.

### Materials

Stimuli were presented on an Iiyama MM904UTA Vision Master Pro 455 cathode ray tube screen with a refresh rate of 85Hz and a resolution of 1280×1024 using a MacPro. A Cambridge Research Systems ColorCal MK II colorimeter was used to calibrate screen luminance, and an accurate pixel per centimetre conversion was obtained by measuring lines on the screen by hand. The screen was viewed through a four-mirror stereoscope (because the set-up was also used for concurrent binocular vision experiments). However, here, the right and left eyes views were always identical (zero binocular disparity). Including the distances between the stereoscope mirrors, the screen was viewed from a distance of 97cm.

### Experiment Design

#### Stimulus Design

All stimuli were viewed from a distance of 97cm created using MATLAB R2014b (The MathWorks Inc, Natick, MA, USA) and Psychtoolbox 3 (Brainard, 1997; Kleiner, Brainard, & Pelli, 2007; Pelli, 1997). Stimuli were presented on a grey background with a luminance of 29.9 cd/m^2^. The stimulus used in the main experimental conditions consisted of a pair of white horizontal lines that were 6.96 degrees long and had a luminance of 59.9cd/m^2^ (see Figure 1). The horizontal lines in the stimulus expanded away from one another to deliver the looming cue. Sekuler (1992) suggests that looming is encoded by the pooling of unidirectional motion signals. Each of the two horizontal lines in the stimulus can be considered an independent set of pooled unidirectional motion signals. The two lines have opposing motion directions, generating a looming signal. Horizontal lines were used as they create no horizontal disparity, so that there would be no conflict between the looming cues and the lack of changing binocular disparity. The pair of lines simulates a very wide object approaching the observer (whose left and right edges are outside the field of view). Each line remained a constant 0.95 arcmin wide on the screen. The separation between the lines varied over time to simulate motion towards the observer, but at the plane of fixation (the screen distance of 97cm) the separation between the two lines was 2cm. The starting separation of the two lines was therefore 2cm for the speed discrimination conditions where line movement began at the plane of fixation, and 1.66 cm for the speed change discrimination condition where line movement began from 20cm behind the plane of fixation.

**Figure 1.**
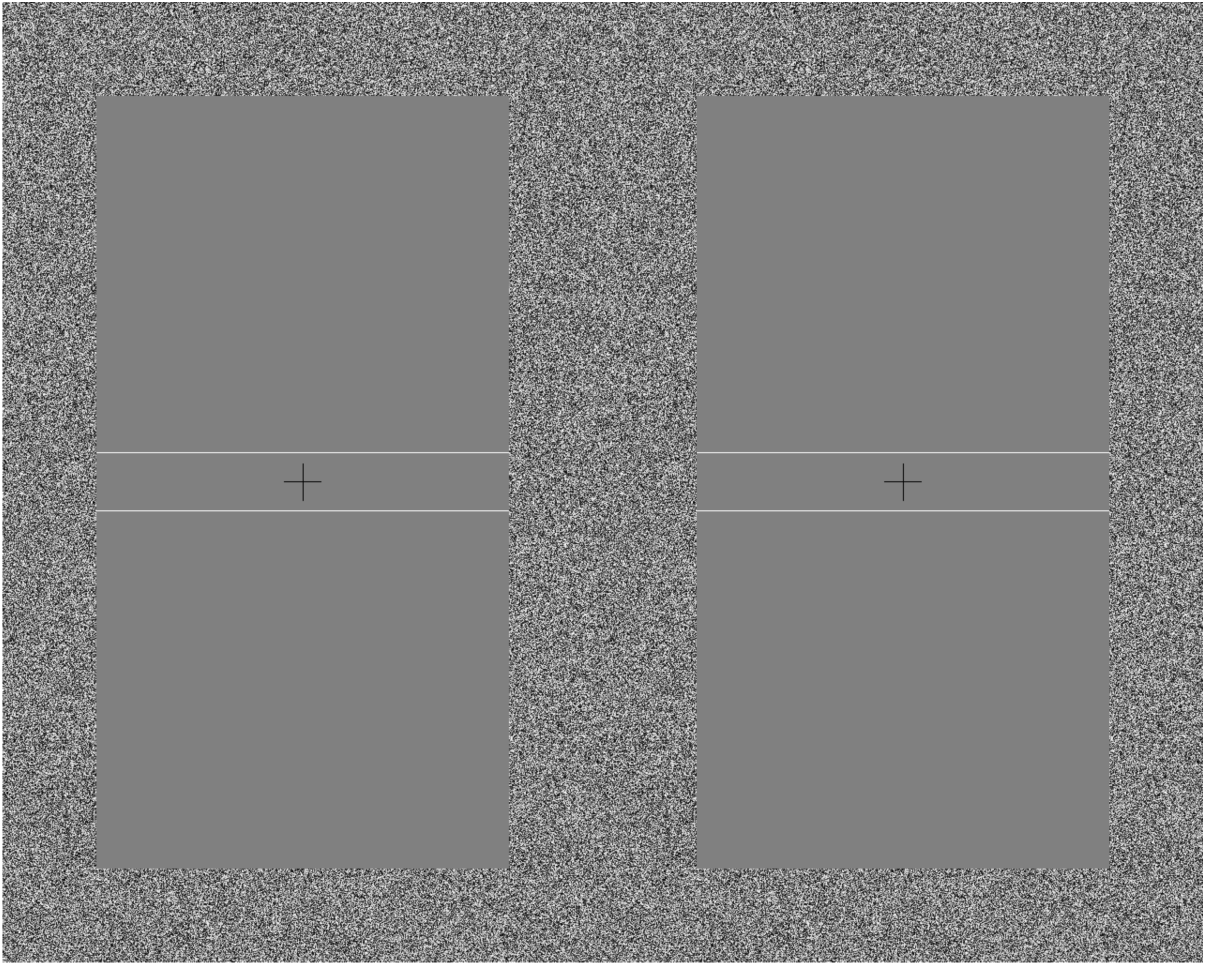
The looming stimulus used for all experimental conditions. The bottom lines moved downwards and the top lines moved upwards to simulate looming motion towards the observer. The left and right halves of the screen were delivered to each eye separately via a 4-mirror stereoscope to deliver a fused percept, but there was no binocular disparity displayed in the stimulus.

Stimuli were presented with a central black fixation cross with a luminance of 0.09 cd/m^2^. This was 37.9 long by 37.9 arcmin wide. To indicate when a response was required, a black by 56.9 arcmin box of the same luminance appeared around the fixation cross. The fixation cross and box had line widths of 0.95 arcmin. Throughout each trial, an aperture frame of approximately uniformly distributed luminance noise, with individual pixels randomly assigned grey levels, was displayed around all stimuli. The aperture frame was 1.58 degrees wide, had a minimum luminance of 0.09 cd/m^2^ and a maximum luminance of 59.9 cd/m^2^. The half-screen visible through the stereoscope used was 10.1 degrees wide and 16.2 degrees tall. Within the aperture frame a rectangle 6.96 degrees wide and 13.0 degrees tall was used for stimulus presentation.

#### Main Experiment Conditions

The stimulus used contained constant world speed (accelerating retinal speed) for three main experimental conditions:

I. *Speed change discrimination* containing no useful distance or duration information. The task was to judge which interval contained a speed change.
II. *Duration (speed discrimination*), containing duration and speed information. The task was to discriminate which interval contained faster motion.
III. *Distance (speed discrimination)* containing distance and speed information. The task was to again discriminate which interval contained faster motion.

#### Determining which cue is used for speed discrimination with catch trials

For each of the two speed discrimination conditions (*Distance* and *Duration*), we included a total of 30 ‘catch trials’ designed to reveal what cues participants were using to perform the task. These trials contained standard and catch *intervals*. The catch intervals contained the same speed as the standard interval but had increased distance and duration (something travelling for longer at the same speed travels further). The duration of the catch interval was always 1.070s, and the standard interval duration was the same as for the main experimental trials in that block (0.717s for *Duration* and 0.506 for *Distance*). This difference results in catch intervals that should be easily discriminable if the participants are using the distance and/or duration cues.

We coded a participant’s response as “correct” if they chose the catch interval and “incorrect” if they chose the standard. Because the catch trials contained the same speed in both intervals, if participants used speed only we would expect performance to be at 50%. However, the catch intervals contained increased distance travelled and duration compared to the standard. Increased distance travelled might be associated with an object appearing to travel faster. If so, participants should choose the catch interval more often. An increased duration might be associated with an object appearing to travel slower. If using duration, participants would choose the standard interval more often. No matter what rule the participant used, if either distance or duration was used in addition to, or instead of speed to perform the task, we would expect performance to be different from 50% for these catch trials.

We made a simple assumption: that participants would use only one cue (duration, distance or speed), and they would attempt to use the same cue across all trials. Here, we had a null hypothesis that if people are using solely speed information, they would be picking the catch trial 50% of the time, because the catch and the standard intervals contain the same speed. We can use the binomial test to determine if we can reject the null hypothesis. To do this we use the binomial distribution to find which values were outside of the 95% confidence interval for the null hypothesis. Values outside of the confidence intervals indicate that we reject the null hypothesis, and suggest that participants were using a cue other than speed to complete the task. For our 30 catch trials per condition, the 95% binomial proportion confidence interval for 50% performance is between 30% and 70%. Thus, if people picked the catch 21 or more times (70% of occasions or more), we can reject the null hypothesis and infer that participants were not only using speed. If the catch interval was picked 9 or less times (30% of occasions or less), participants were again not only using speed.

As the distance and duration were greater in the catch interval, we could infer which cue participants were using to complete the speed discrimination task. We can do this if we assume participants are using a rule that is consistent with speed judgments.

*Speed*: If participants used only speed, not duration or distance, to make their judgement, they would pick the catch interval on 50% of occasions (as we used a forced-choice task and the speeds in each interval were identical). In this scenario we would accept the null hypothesis and performance would be consistent with participants using only speed information.

*Distance*: If participants used the distance cue to make their judgement, they would pick the catch interval significantly more than 50% of the time (because the distance travelled in the catch interval was further, and something that travels further may be thought of as travelling faster). In this scenario we would reject the null hypothesis that the participant is using only speed information.

*Duration*: If participants used the duration cue to make their judgement, they would pick the catch interval significantly less than 50% of the time (because the standard interval had the shorter duration, and something that travels the same distance in a shorter duration may be thought of as travelling faster). Again in this scenario we would reject the null hypothesis that the participant is using only speed information.

#### Training

Two further stimuli were used only for training purposes before the main experiment began: Training I: a square drifting grating with a spatial frequency of 1 cycle per degree which moved from left to right at a constant retinal speed. The grating was 4 degrees tall and 4 degrees wide. Training II: a pair of white vertical lines with a luminance of 59.9 cd/m^2^ that moved from left to right at a constant retinal speed. These lines were each 13.0 degrees tall and 0.95 arcmin wide. All the code used in this experiment is available online at https://osf.io/xvs5n/.

### Procedure

#### Speed Change Discrimination

For the speed change discrimination condition, participants completed a 2-interval forced choice task, with a 7-level method of constant stimuli design. One interval contained an instantaneous speed change from a slower to a faster speed, the other contained motion at a constant speed. Participants were asked to identify the interval that contained the speed change. Each interval began with the stimulus appearing and remaining stationary for 250ms, before moving for 1 second. If the interval contained a change in speed, it occurred after 500ms of motion. There was a gap of 1s between the two intervals. Before the first interval participants heard one beep; before the second they heard two beeps. A run consisted of 210 trials divided into three blocks, each with 10 trials per level, giving 70 trials per block. These blocks were presented in a random order amongst those for a different study, with other speed change blocks containing either binocular or binocular and looming cues to motion in depth.

A standard speed of 40 cm/s towards the observer was used, which translated to a speed range of 20.1-46.3 arcmin/s on the retina over the full interval. In each successive level of the condition, the speed before the speed change decreased by 5 cm/s, and the speed after the change increased by 5 cm/s in the world. This meant that the test speeds were (speed before change-speed after change): 40cm/s-40cm/s, 35cm/s-45cm/s, 30cm/s-50cm/s, 25cm/s-55cm/s, 20cm/s-60cm/s, 15cm/s-65cm/s, and 10cm/s-70cm/s. In the maximum speed change level, from 10cm/s-70cm/s, this translates into a range of retinal speeds between 5.02-5.48 arcmin/s for the period before the instantaneous speed change, and a range of retinal speeds between 38.4-80.9 arcmin/s after the change.

All participants were required to complete three training blocks prior to the main experiment, which used the same task, but different stimuli. The Training I stimulus, a drifting grating, was used first in a speed change discrimination task like that used in the main experiment, and participants received audio feedback on their responses. The second block was identical, except participants no longer received feedback. The third block used the Training II stimulus (two vertical lines) in a speed change discrimination task with no feedback. Stimuli in these 3 training blocks all used constant retinal speed and contained the same two speed change levels. Participants compared a standard stimulus moving at 283.4 arcmin/s to a step speed change from 212.6 to 354.1 arcmin/s, or from 70.9 to 495.3 arcmin/s. The purpose of the training blocks was to introduce participants to the task by first using a commonly-used motion stimulus undergoing lateral motion (Training I), then a stimulus more similar to that used in the main experiment (Training II).

#### Speed Discrimination with Duration or Distance

For the speed discrimination conditions containing either duration or distance cues (*Duration* and *Distance*), participants viewed two temporal intervals both containing stimuli travelling at a constant world speed. The task was to pick the interval in which the stimulus moved faster. Participants were told to judge the speed, but were not given any instructions about use of distance or duration information. For each condition two consistent cues were available (either speed and duration, or speed and distance). Our aim was to test whether performance was better for one condition or the other and compare these speed discrimination conditions to the *Speed Change* condition (where only speed could be used). A 7-level method of constant stimuli design was used. Each interval began with the stimulus appearing and remaining stationary for 0.250s. The *Distance* condition, where duration was fixed but distance information was available, contained 0.506s of motion in both the standard and test intervals. The distances presented ranged between 20 cm in depth in the standard speed level and 35 cm in depth in the maximum speed level and 42.8 cm in the catch interval. The *Duration* condition, where distance was fixed but duration information was available, consisted of a test interval containing a range of durations to keep the distance travelled constant at 28.7cm towards the observer. The durations ranged between 0.412s of motion for the maximum speed level, and 0.717s of motion for the minimum speed level. The standard interval in the *Duration* condition was presented for 0.717s. There was a gap of 1s between intervals, and before the first interval participants heard one beep; before the second they heard two beeps. The duration of the catch interval was always 1.070s, and the standard interval duration was the same as for the main experimental trials in that block (0.717s for *Duration* and 0.506 for *Distance*). The *Duration* and *Distance* conditions each consisted of 210 main experiment trials, which were randomly interleaved with the catch trials presented in 3 blocks with 10 trials per level, to provide participants with breaks. Blocks contained either only *Duration* or only *Distance* trials, but blocks of each condition were presented in a random order.

In both speed discrimination conditions, the standard stimulus had a speed of 40 cm/s in the world. Successive test levels increased in speed by 5 cm/s, to a maximum speed of 70 cm/s. This meant that test levels had speeds of: 40cm/s, 45cm/s, 50cm/s, 55cm/s, 60cm/s, 65cm/s and 70cm/s. We chose these speeds so that they would be matched to the second speeds used in the test intervals of the speed change discrimination condition (see above). In the *Distance* conditions, these translated into retinal speeds of 29.2 – 46.3 arcmin/s in the standard level, and 51.2-125.0 arcmin/s in the maximum speed level. In the *Duration* conditions, these world speeds translated into retinal speeds of 29.2-58.8 arcmin/s in the standard level, and 51.2-103.7 arcmin/s in the maximum speed level. In the catch trials, the speed of stimuli in the catch and standard intervals was 40 cm/s in the world, which translates differently into accelerating retinal speeds depending on the duration of the interval. In the *Duration* condition, the standard interval speeds ranged between 29.2 – 58.8 arcmin/s, and the catch interval speeds ranged between 29.2 – 93.4 arcmin/s. In the *Distance* condition, the speeds in the standard interval ranged between 29.2 – 46.6 arcmin/s, while the catch interval speeds ranged between 29.2 – 93.4 arcmin/s.

Prior to the experiment start, participants completed two speed discrimination training blocks, one *Duration* and one *Distance*, using the horizontal line stimulus used in the main experiment. Participants received audio feedback on their responses and two levels of speed difference were used. The standard stimulus moved towards the observer at 40 cm/s, whilst the test stimulus had a speed of 50 cm/s or 70 cm/s in the world.

### Analysis

We measured 75% thresholds for speed change discrimination and speed discrimination by fitting cumulative normal psychometric functions using MATLAB R2014b (The MathWorks Inc, Natick, MA, USA) and the Palamedes toolbox (Prins & Kingdom, 2009). With this we found the speed change required to respond correctly on 75%of occasions, or the difference in speed required to respond correctly on 75% of occasions, both as a function of world speed (cm/s). This method was the most appropriate method of analysing the data from the speed change discrimination and speed discrimination tasks to avoid bias. We also considered analysing our data in terms of the proportion speed difference or speed change, as we have done previously (see Lee et al., 2019), and in terms of the arcmin/s difference or change in speed. However, these methods introduced bias to the results because the arcmin/s changes in speed were not identical between conditions, and consequently the proportion speed changes and differences that were based on these arcmin/s values were also not identical between conditions. Example psychometric functions are shown in Figure 2. All data, experimental code and analysis code is available online at https://osf.io/xvs5n/.

**Figure 2.**
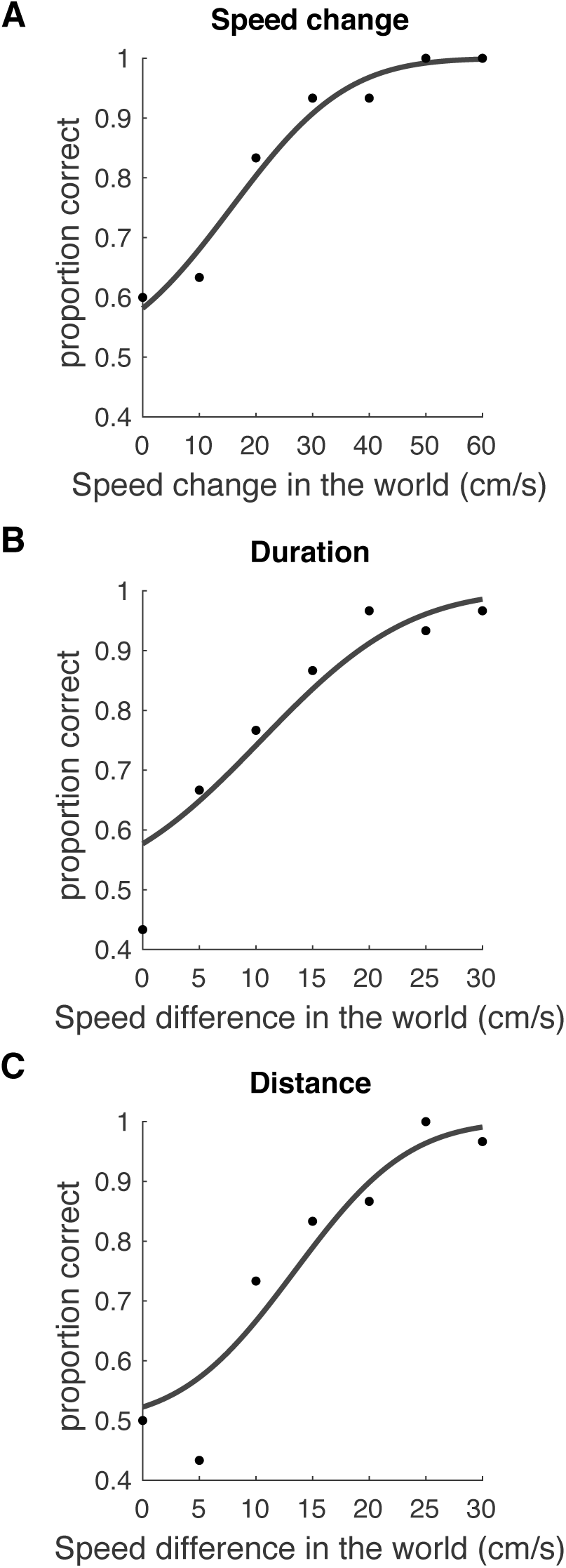
Example psychometric functions in the speed change discrimination condition (A), the *Duration* speed discrimination condition (B), and the *Distance* speed discrimination condition (C). (B) and (C) are data from the same participant.

JASP (Version 0.9.1; JASP Team, 2018) was used to conduct a pair of two-sample t-tests. Threshold values between the *Speed Change* discrimination condition and the *Duration* speed discrimination condition, and the *Speed Change* discrimination condition and the *Distance* speed discrimination condition were compared. This was done to determine if there was a difference in threshold between the speed change discrimination condition and the two speed discrimination conditions. A paired t-test was then used to compare thresholds between the *Distance* and *Duration* speed discrimination conditions, to investigate if thresholds varied depending on whether duration or distance information were available respectively. The significance level for all t-tests was taken as 0.0167 (Bonferroni correction for 3 comparisons). We also measured Pearson’s correlation coefficient between the catch trial results for each participant in each speed discrimination condition to observe whether individual participants consistently used the same cue to complete the catch trials in both the *Duration* and *Distance* conditions.

## Results

### Comparing speed discrimination and speed change discrimination

Speed change discrimination was more difficult than speed discrimination when duration or distance information was available (Figure 3). *Speed change discrimination* thresholds were significantly higher than those for *speed discrimination* with a fixed distance and variable duration (*Duration* condition; t(16) = 4.180, p < 0.001) and also were higher than for *speed discrimination* with a fixed duration and variable distance (*Distance* condition; t(16) = 3.009, p = 0.008). There was no significant difference in thresholds between the *Duration* and *Distance* conditions (t(6) = −0.381, p = 0.716). It appears that speed change discrimination is a more difficult task than speed discrimination irrespective of the cues available to distinguish between the speeds.

**Figure 3.**
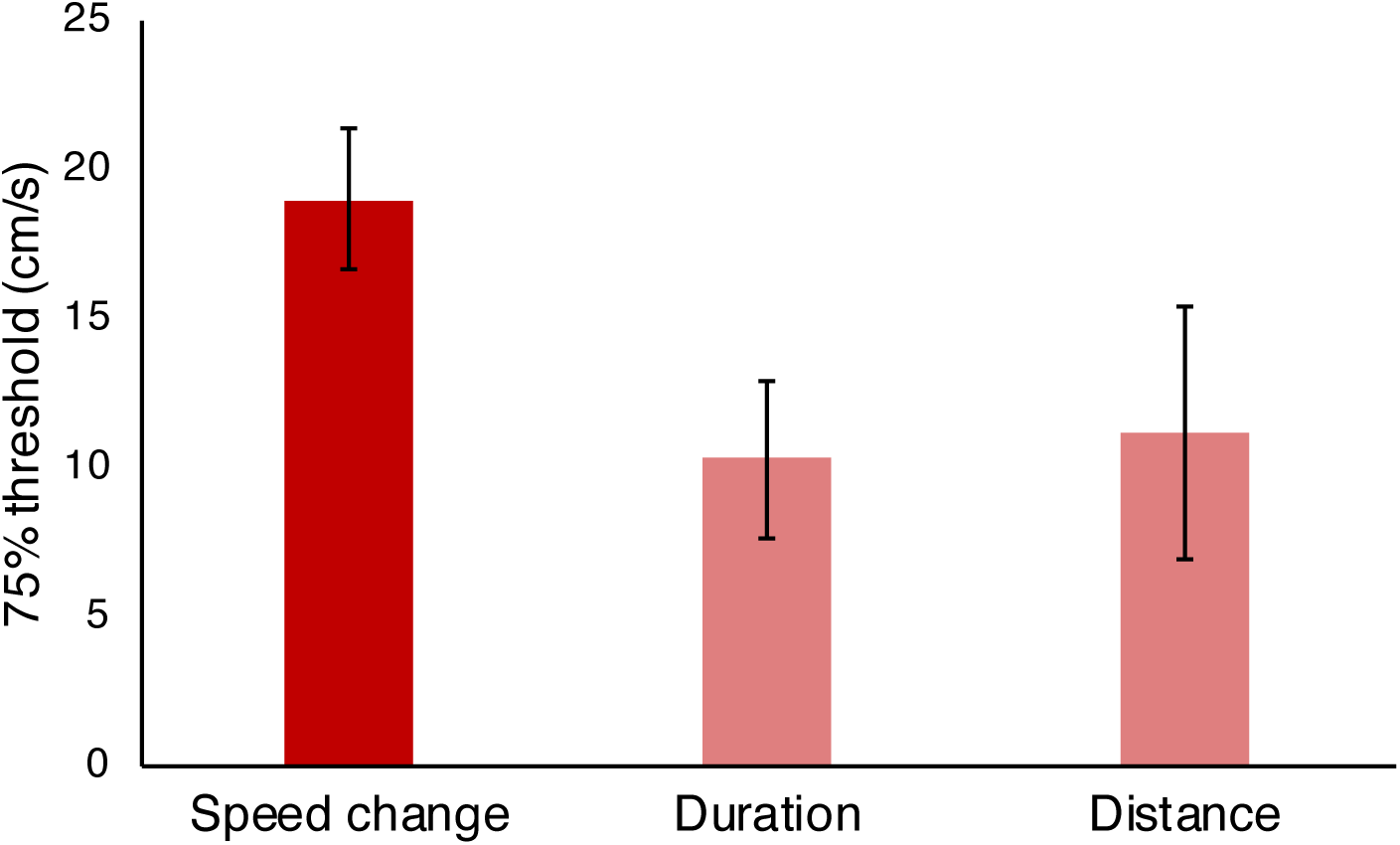
75% thresholds for the *Speed change* discrimination condition (n = 11) and the *Duration* and *Distance* speed discrimination conditions, both n = 7. Error bars are 95%confidence intervals.

### Cue usage in catch trials

Our catch trial analysis showed that participants used a variety of strategies to complete the task (Figure 4). We categorised the catch trial results into 3 groups based on binomial proportion confidence intervals:

**Figure 4.**
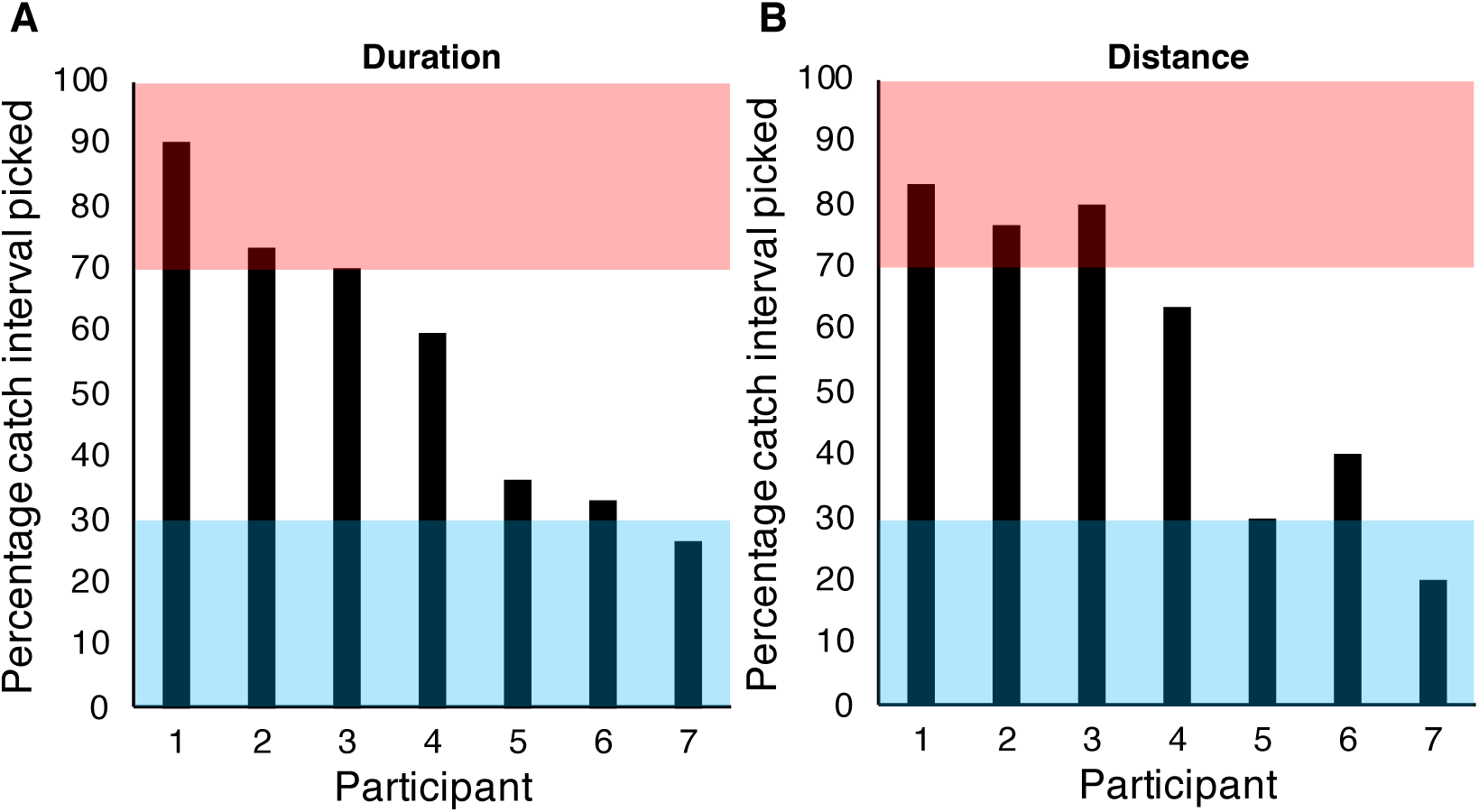
The percentage of occasions the catch interval was picked over the standard interval in the catch trials for (A), the *Duration* speed discrimination condition and (B), the *Distance* speed discrimination condition. Participants whose data lie in the pink region used distance cues. Within the blue region, participants used duration information. Data in the white region suggest participants used speed.

I. 0-30% catch interval picked – we reject the null hypothesis that participants used only speed. Participant was using duration as a cue.
II. 70-100% catch interval picked – we reject the null hypothesis that participants used only speed. Participant was using distance as a cue.
III. 31-69% catch interval picked – we do not reject the null hypothesis. Participant was primarily using speed to make their judgements in the catch trials.

From the pattern of data in Figure 4A, it appeared that in the *Duration* speed discrimination condition, 1 participant used the shorter duration information, 3 participants used speed information and 3 participants used the longer distance information to complete the task. In the *Distance* speed discrimination condition, 2 participants used the shorter duration, 2 participants used speed information and 3 participants used the longer distance information to complete the task (see Figure 4B). Our data suggest that different participants used different cues to complete speed discrimination tasks.

### Individual participant data

Given that some individual participants appear to use cues other than speed in the catch trials, we might expect that individual participants may perform better in the main experiment condition where they could use the cue they favoured in the catch trials. For example, we would expect that Participant 1 in Figure 4, who appeared to use distance information to make their catch trial judgements, would have improved performance in the *Distance* condition of the main experiment, because they could use the cue their favoured cue from the catch trials. In the *Duration* condition of the main experiment, we would expect worse performance from Participant 1, because there was no distance information available for them to use.

There were marked individual differences in performance, as has been found previously in experiments involving speed discrimination that have used naїve observers (Manning, Thomas, & Braddick, 2018). However, individual participants did not show a pattern of thresholds in the main experiment that would be predicted from their behaviour during the catch trials. Figure 5 illustrates that there was very little difference in threshold between the two speed discrimination conditions in the main experiment, for all participants.

**Figure 5.**
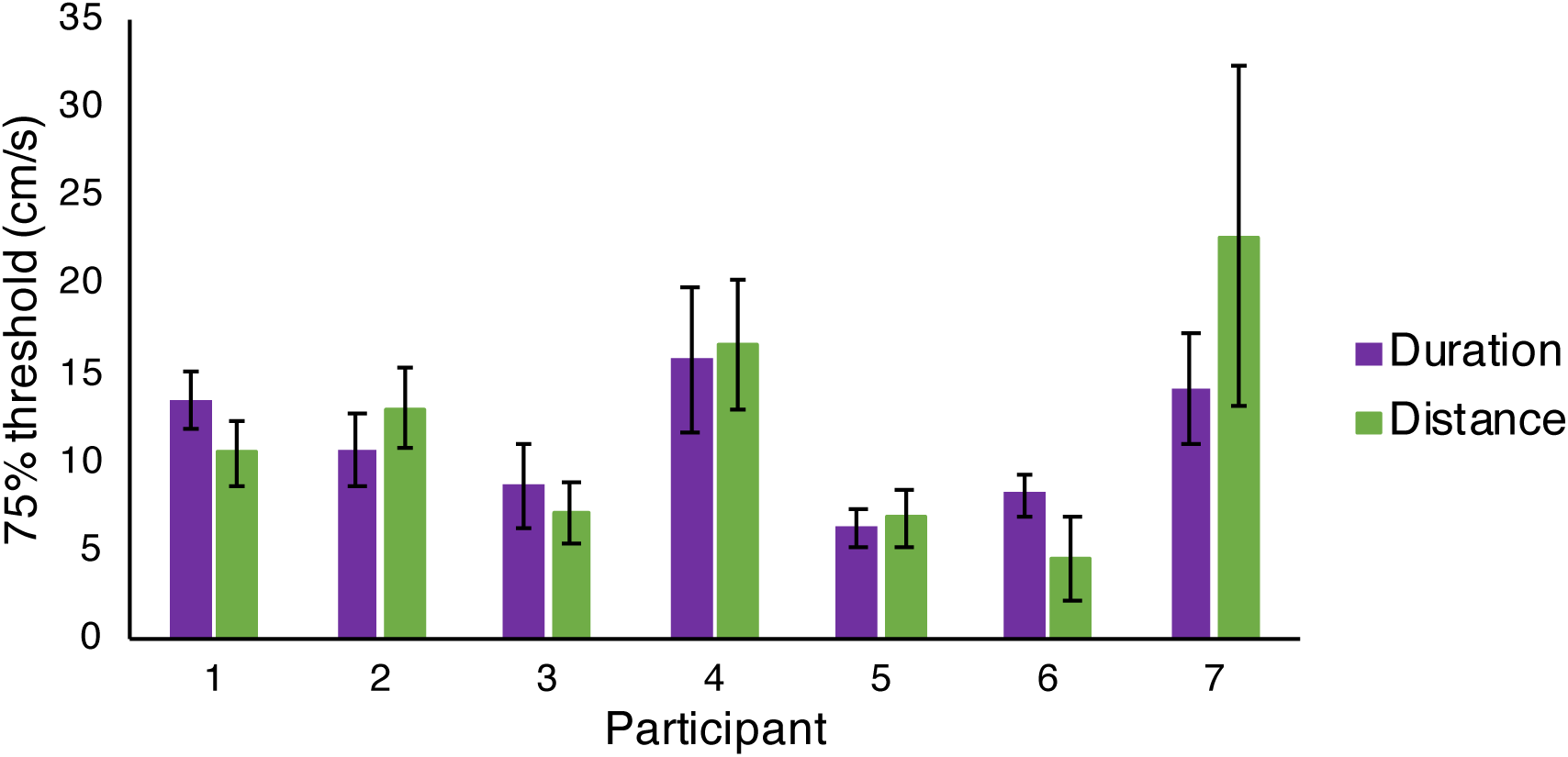
Individual 75%thresholds for each participant in the two speed discrimination conditions (*Duration* and *Distance*, n = 7). Error bars are standard error of the mean.

### Do participants use the same cues in the catch and main experiment trials?

Notice that individual participants were consistent in their choice of catch trial cue usage between the two catch conditions (figure 4). A significant strong correlation of percentage catch interval picked per participant was found between the two catch conditions (r = 0.963, n = 7, p < 0.001). This suggests participants may have been using cues to perform the task in the catch trials that would not be helpful if used in the main experimental trials. For example, 3 participants appeared to use distance information in the catch trials that were included as part of the *Duration* speed discrimination condition (see Figure 4A). In this *Duration* condition, speed and duration cues were available but the distance travelled was fixed. This means those participants used distance information in the catch trials despite it being impossible to successfully use distance to make judgements in the main experiment trials of this condition. As the catch trials were interleaved with the main experimental trials, this suggests that participants may have been changing between using different cues from one trial to the next when completing the speed discrimination task.

## Discussion

The aims of this work were to first determine if speed change discrimination for looming stimuli is a more difficult task than speed discrimination, and second, to investigate if participants use distance or duration information, instead of speed information, when it is available in speed discrimination tasks. If so, this might explain the apparent difficulty of speed change discrimination judgements. To do this we measured discrimination thresholds for the two different tasks and included manipulations of the speed discrimination stimuli in the form of ‘catch trials’, which were designed to reveal the use of distance, duration, or speed information.

We found speed change discrimination to be significantly more difficult than speed discrimination for looming stimuli. This result is in agreement with the previous literature that has studied at speed change discrimination for 2D motion and other varieties of 3D motion (Lee et al., 2019; Monen & Brenner, 1994; Snowden & Braddick, 1991). The majority of our individual participants had 75% thresholds below 20 cm/s in the speed discrimination conditions (*Duration* and *Distance*, see Figure 5).

In experiments from other labs using a different type of speed change discrimination task, where a participant determines whether a stimulus has increased or decreased in speed, the distinction between speed change and speed discrimination thresholds is slightly less clear. Thresholds have been reported to be roughly 3 times higher when two speeds are presented consecutively than when there was a period of 1 second between the two speeds, but only when only when the duration of motion was short (Mateeff et al., 2000). In another study thresholds as low as 12% have been found for this type of experiment (Hick, 1950). However, unlike in the two-interval forced choice speed change discrimination task used in our study, Hick’s (1950) experimental setup left it clear when a speed change had occurred, which could have made their task easier and explain their lower thresholds.

Our manipulations of the catch trials in the speed discrimination task suggested that participants were not exclusively using speed to complete the task. We found some participants who appeared to use speed, some duration, and some distance in the catch trials, and the cue use correlated between the catch trials in the *Duration* and *Distance* conditions. This means in some cases participants would have used catch trial strategies that they could not have used successfully for other trials in the main experiment. For example, in Figure 4, participant 1 appears to often be using the distance cue to make their judgements in the catch trials in both the *Duration* condition (Figure 4A) and the *Distance* condition (Figure 4B). However, participant 1 would not have been able to successfully use distance to perform the speed discrimination task in the main experiment trials of the *Duration* condition: because the distance cue was not informative. As participant 1 was able to perform this task well (see Figure 5), we can infer that participants may have been able to use multiple cues to complete the speed discrimination task, and, as the catch trials were interleaved amongst the other experimental trials, that participants could have been switching between using different cues from trial to trial. It is also possible that participants may have used a combination of differently-weighted cues to make their judgements in each trial, but our design was not aimed at testing this hypothesis. As such, we cannot address this possibility here.

The finding that participants may have used cues other than speed to complete the speed discrimination task is supported by other studies that have shown that participants may not use speed information in isolation to discriminate speed (Mandriota, Mintz, & Notterman, 1962; Smith & Edgar, 1991). Mandriota et al. (1962) found that adding distance and duration cues improved speed discrimination performance, whilst Smith and Edgar (1991) reported that discrimination of speed and temporal frequency were inseparable when using drifting grating stimuli, suggesting speed may not be used in isolation.

On the other hand, our catch trial findings conflict with research suggesting participants use speed in speed discrimination tasks with both 2D and 3D motion (Harris & Watamaniuk, 1996; Lappin, Bell, Harm, & Kottas, 1975; McKee, 1981; McKee et al., 1986; Orban et al., 1984; Pasternak, 1987). However, of these studies, only Lappin et al. (1975) included more than 3 human participants in their study, although Pasternak (1987) also had 9 cats as subjects. In our study all participants were fully naїve and inexperienced observers. Three of these previous studies had at least one author as a participant who would have been aware of the aims of the experiment (Harris & Watamaniuk, 1996; McKee et al., 1986; Pasternak, 1987). It is possible that if a larger number of naїve participants had been used in these experiments, a greater range of cue usage in speed discrimination tasks may have been demonstrated.

If participants do not always use speed in speed discrimination tasks, as suggested here, this supports the idea that speed change discrimination may be more difficult than speed discrimination because participants cannot use distance or duration cues in the latter task. However, there are other possible explanations for why speed change discrimination is a difficult task. For example, it is possible that participants may combine or integrate speed information over space or time in such a way that it makes speed change discrimination tasks where two speeds are presented consecutively more difficult. This would be an interesting avenue for future research.

## Acknowledgements

This work was supported by the Biotechnology and Biological Sciences Research Council (BBSRC; https://bbsrc.ukri.org/) [grant number BB/M010996/1 to ARIL, BB/N018516/1 to JMA and BB/M001660/1 to JMH]. The funders had no role in study design, data collection and analysis, decision to publish, or preparation of the manuscript.

## References

Beverley, K. I., & Regan, D. (1979). Seperable aftereffects of changing-size and motion-in-depth: different neural mechanisms? Vision Research, 19, 727–723. https://doi.org/ https://doi.org/10.1016/0042-6989(79)90251-7

Brainard, D. H. (1997). The Psychophysics Toolbox. Spatial Vision, 10(4), 433–436. https://doi.org/10.1163/156856897X00357

Brooks, K. R. (2002). Interocular velocity difference contributes to stereomotion speed perception. Journal of Vision, 2(3), 218–231. https://doi.org/10.1167/2.3.2

Brooks, K. R., & Mather, G. (2000). Perceived speed of motion in depth is reduced in the periphery. Vision Research, 40(25), 3507–3516. https://doi.org/10.1016/S0042-6989(00)00095-X

Brooks, K. R., & Stone, L. S. (2004). Stereomotion speed perception: Contributions from both changing disparity and interocular velocity difference over a range of relative disparities. Journal of Vision, 4(12), 1061–1079. https://doi.org/10.1167/4.12.6

Brooks, K. R., & Stone, L. S. (2006). Stereomotion suppression and the perception of speed: accuracy and precision as a function of 3D trajectory. Journal of Vision, 6(11), 1214–1223. https://doi.org/10.1167/6.11.6

de Bruyn, B., & Orban, G. A. (1988). Human velocity and direction discrimination measured with random dot patterns. Vision Research, 28(12), 1323–1335. https://doi.org/10.1016/0042-6989(88)90064-8

Harris, J. M., & Watamaniuk, S. N. J. (1995). Speed discrimination of motion-in-depth using binocular cues. Vision Research, 35(7), 885–896. https://doi.org/10.1016/0042-6989(94)00194-Q

Harris, J. M., & Watamaniuk, S. N. J. (1996). Poor speed discrimination suggests that there is no specialized speed mechanism for cyclopean motion. Vision Research, 36(14), 2149–2157. https://doi.org/10.1016/0042-6989(95)00278-2

Heidenreich, S. M., & Turano, K. A. (1996). Speed discrimination under stabilized and normal viewing conditions. Vision Research, 36(12), 1819–1825. https://doi.org/10.1016/0042-6989(95)00270-7

Hick, W. E. (1950). The threshold for sudden changes in the velocity of a seen object. Quarterly Journal of Experimental Psychology, 2(September), 33–41. https://doi.org/10.1080/17470215008416572

Kleiner, M., Brainard, D. H., & Pelli, D. (2007). What’s new in Psychtoolbox-3? Perception, 36 (ECVP Abstract Supplement). Retrieved from http://psychtoolbox.org/credits/

Lappin, J. S., Bell, H. H., Harm, O. J., & Kottas, B. (1975). On the relation between time and space in the visual discrimination of velocity. Journal of Experimental Psychology. Human Perception and Performance, 1(4), 383–394. https://doi.org/10.1037/0096-1523.1.4.383

Lee, A. R. I., Ales, J. M., & Harris, J. M. (2019). Speed change discrimination for motion in depth with constant world and retinal speeds. PLoS ONE, 14(4), e0214766. https://doi.org/10.1371/journal.pone.0214766

Mandriota, F. J., Mintz, D. E., & Notterman, J. M. (1962). Visual Velocity Discrimination: Effects of Spatial and Temporal Cues. Science, 138(3538), 437–438.

Manning, C., Thomas, R. T., & Braddick, O. J. (2018). Can speed be judged independent of direction? Journal of Vision, 18(6), 1–16. https://doi.org/10.1167/18.6.15

Mateeff, S., Dimitrov, G., Genova, B., Likova, L., Stefanova, M., & Hohnsbein, J. (2000). The discrimination of abrupt changes in speed and direction of visual motion. Vision Research, 40(4), 409–415. https://doi.org/10.1016/S0042-6989(99)00185-6

McKee, S. P. (1981). A local mechanism for differential velocity detection. Vision Research, 21(4), 491–500. https://doi.org/10.1016/0042-6989(81)90095-X

McKee, S. P., Silverman, G. H., & Nakayama, K. (1986). Precise velocity discrimination despite random variations in temporal frequency and contrast. Vision Research, 26(4), 609–619.

McKee, S. P., & Watamaniuk, S. N. J. (1994). The psychophysics of motion perception. In A. T. Smith & R. J. Snowden (Eds.), Visual Detection of Motion (pp. 85–114). London: Academic Press.

McKee, S. P., & Welch, L. (1985). Sequential recruitment in the discrimination of velocity. Journal of the Optical Society of America A, 2(2), 243–251. https://doi.org/10.1364/JOSAA.2.000243

Monen, J., & Brenner, E. (1994). Detecting changes in one’s own velocity from the optic flow. Perception, 23(6), 681–690. https://doi.org/10.1068/p230681

Orban, G. A., de Wolf, J., & Maes, H. (1984). Factors influencing velocity coding in the human visual system. Vision Research, 24(1), 33–39. https://doi.org/10.1016/0042-6989(84)90141-X

Pasternak, T. (1987). Discrimination of differences in speed and flicker rate depends on directionally selective mechanisms. Vision Research, 27(11), 1881–1890. https://doi.org/10.1016/0042-6989(87)90054-X

Pelli, D. G. (1997). The VideoToolbox software for visual psychophysics: transforming numbers into moview. Spatial Vision, 10(4), 437–442. https://doi.org/10.1163/156856897X00366

Prins, N., & Kingdom, F. A. A. (2009). Palamedes: Matlab routines for analyzing psychophysical data. Retrieved from http://www.palamedestoolbox.org/

Regan, D., & Beverley, K. I. (1978). Illusory motion in depth: Aftereffect of adaptation to changing size. Vision Research, 18, 209–212. https://doi.org/10.1016/0042-6989(78)90188-8

Sekuler, A. B. (1992). Simple-pooling of unidirectional motion predicts speed discrimination for looming stimuli. Vision Research, 32(12), 2277–2288. https://doi.org/10.1016/0042-6989(92)90091-V

Smith, A. T., & Edgar, G. K. (1991). The separability of temporal frequency and velocity. Vision Research, 31(2), 321–326. https://doi.org/10.1016/0042-6989(91)90121-K

Snowden, R. J., & Braddick, O. J. (1991). The temporal integration and resolution of velocity signals. Vision Research, 31(5), 907–914. https://doi.org/10.1016/0042-6989(91)90156-Y

Wardle, S., & Alais, D. (2013). Evidence for speed sensitivity to motion in depth from binocular cues. Journal of Vision, 13(2013), 1–16. https://doi.org/10.1167/13.1.17.Introduction

